# Oct1/Pou2f1 is selectively required for gut regeneration and regulates gut malignancy

**DOI:** 10.1101/414391

**Authors:** Karina Vázquez-Arreguín, Claire Bensard, John C. Schell, Eric Swanson, Xinjian Chen, Jared Rutter, Dean Tantin

## Abstract

The transcription factor Oct1/Pou2f1 promotes poised gene expression states, mitotic stability, glycolytic metabolism and other characteristics of stem cell potency. To determine the effect of Oct1 loss on stem cell maintenance and malignancy, we deleted Oct1 in two different mouse gut stem cell compartments. Oct1 deletion preserved homeostasis in vivo and the ability to generate cultured organoids in vitro, but blocked the ability to regenerate after treatment with dextran sodium sulfate, and the ability to maintain organoids after passage. In a chemical model of colon cancer, loss of Oct1 in the colon severely restricted tumorigenicity. In contrast, loss of one or both *Oct1* alleles progressively increased tumor burden in a colon cancer model driven by loss of heterozygosity of the tumor suppressor gene *Apc.* The different outcomes are consistent with prior findings that Oct1 promotes mitotic stability, and consistent with different gene expression signatures associated with the two models. These results reveal that Oct1 is selectively required for gut regeneration, and has potent effects in colon malignancy, with outcome (pro-oncogenic or tumor suppressive) dictated by tumor etiology.

**Author summary:** Colorectal cancer is the second leading cause of cancer death in the United States. Approximately 35% of diagnosed patients eventually succumb to disease. The high incidence and mortality due to colon cancer demand a better understanding of factors controlling the physiology and pathophysiology of the gastrointestinal tract. Previously, we and others showed that the widely expressed transcription factor is expressed at higher protein levels in stem cells, including intestinal stem cells. In this study we use a conditional mouse *Oct1* (*Pou2f1*) allele deleted in two different intestinal stem cell compartments. The results indicate that Oct1 loss is dispensable for maintenance of the mouse gut, but required for regeneration. We also tested Oct1 loss in the context of two different mouse colon cancer models. We find that Oct1 loss has opposing effects in the two models, and further that the two models are associated with different gene expression signatures. The differentially expressed genes are enriched for previously identified Oct1 targets, suggesting that differential gene control by Oct1 is one mechanism underlying different outcomes.

## Introduction

Multiple studies identify a correlation between tumor aggressiveness and elevated expression of Oct1/Pou2f1 [1-9], including in the GI tract [10-13].

Oct1 is a widely expressed POU domain transcription factor related to the embryonic stem cell master transcription factor, Oct4 [1,14]. Oct1 promotes glycolytic metabolism and mitotic stability [15-17]. It also promotes poised gene expression states, i.e. the ability of transcriptionally silent target genes to remain inducible following the correct developmental cues [18,19]. Oct1 loss is associated with increased oxidative metabolism, elevated reactive oxygen species, hypersensitivity to oxidative and genotoxic stress, and a modest increase in abnormal mitoses [1,15,16,20,21]. Oct1 loss does not compromise cell viability, affect immortalization by serial passage or reduce growth rates in standard culture. Oct1 is however required for oncogenic transformation in soft agar [15]. In a well-characterized *Tp53* null mouse model, loss of even one Oct1 allele suppresses thymic lymphoma [15]. In a variety of malignancies, Oct1 motifs are enriched in coordinately activated genes [3,22-24]. These results indicate that Oct1 plays important roles in stress responses and tumorigenicity.

We and others have shown that Oct1 promotes somatic and cancer stem cell potency in different systems [1,25,26]. For example, in the blood system Oct1 loss allows for primary hematopoietic engraftment but compromises the capacity for serial transplants [1]. Despite the connection between Oct1, stem cells and oncogenic potential, *Oct1* (*Pou2f1*) mRNA is not elevated in stem cells [27]. Instead, Oct1 is regulated post-translationally, e.g. at the level of DNA binding specificity [17], sub-nuclear localization [16,21,28-30], and protein stability [7,25]. Oct1 protein levels are elevated in mouse and human gastric, small intestine and colon stem cells [1,11,31]. Two Oct1 ubiquitin ligases have been identified, Trim21 and BRCA1 [7,25].

Here, we use a conditional Oct1 allele together with two gut stem cell Cre-drivers to determine the role of this protein in the maintenance and regeneration of normal gut cells, and in gastrointestinal malignancy. The Cre-drivers are tamoxifen inducible to provide both tissue-specific and temporal control over Oct1 deletion. We find that Oct1 is dispensable for colon homeostasis, but required for normal regeneration. In the small intestine, Oct1 deletion from Lgr5^+^ cells in vivo allows for the establishment of cultured gut organoids from isolated crypts, but not their maintenance following passage. In a chemical model of colon tumorigenesis, Oct1 loss greatly reduces tumor incidence and size. The tumors that do emerge in this model have escaped Oct1 deletion, indicating a requirement for Oct1. In contrast, a model of colon cancer driven by loss of heterozygosity (LOH) shows a progressive increase in tumor number, though not of grade, as one or both *Oct1* alleles are lost. The increase in tumor initiation indicates a tumor-suppressive role of Oct1 in this context, and is consistent with prior findings that Oct1 promotes mitotic stability. Comparison of gene expression signatures in the two models identifies a set of genes correlating with the model of origin, indicating that the two models have distinct molecular features. This set enriched in direct Oct1 targets.

## Results

### Oct1 is required for colon epithelium regeneration but not homeostasis

We crossed an *Oct1* (*Pou2f1*) conditional allele [18] to Lrig1-CreER^T2^ [32], which upon tamoxifen treatment (single injection of 2 mg in corn oil by gavage, see methods) induces Cre activity in stem cells of the colon. 4 wks after treatment, the colon epithelium was superficially normal (Fig 1A), despite near-global deletion of Oct1 (Fig 1B). Oct1 deletion and normal colonic architecture could be maintained for 150 days (not shown). Histological examination of tissue revealed that Oct1 was deleted in >90% of crypts (Fig 1C). In contrast, non-epithelial cells retained staining (Fig 1D, asterisk). Rarely there were crypt structures that escape deletion (red arrow), whose frequency did not change over time (not shown). Crypts lacking Oct1 appeared normal compared to those that had escaped deletion (Fig 1D, right) and compared to Oct1 sufficient mice (Fig 1D, left). These results show that Oct1 loss has no superficial effect on the colon epithelium.

**Fig 1.**
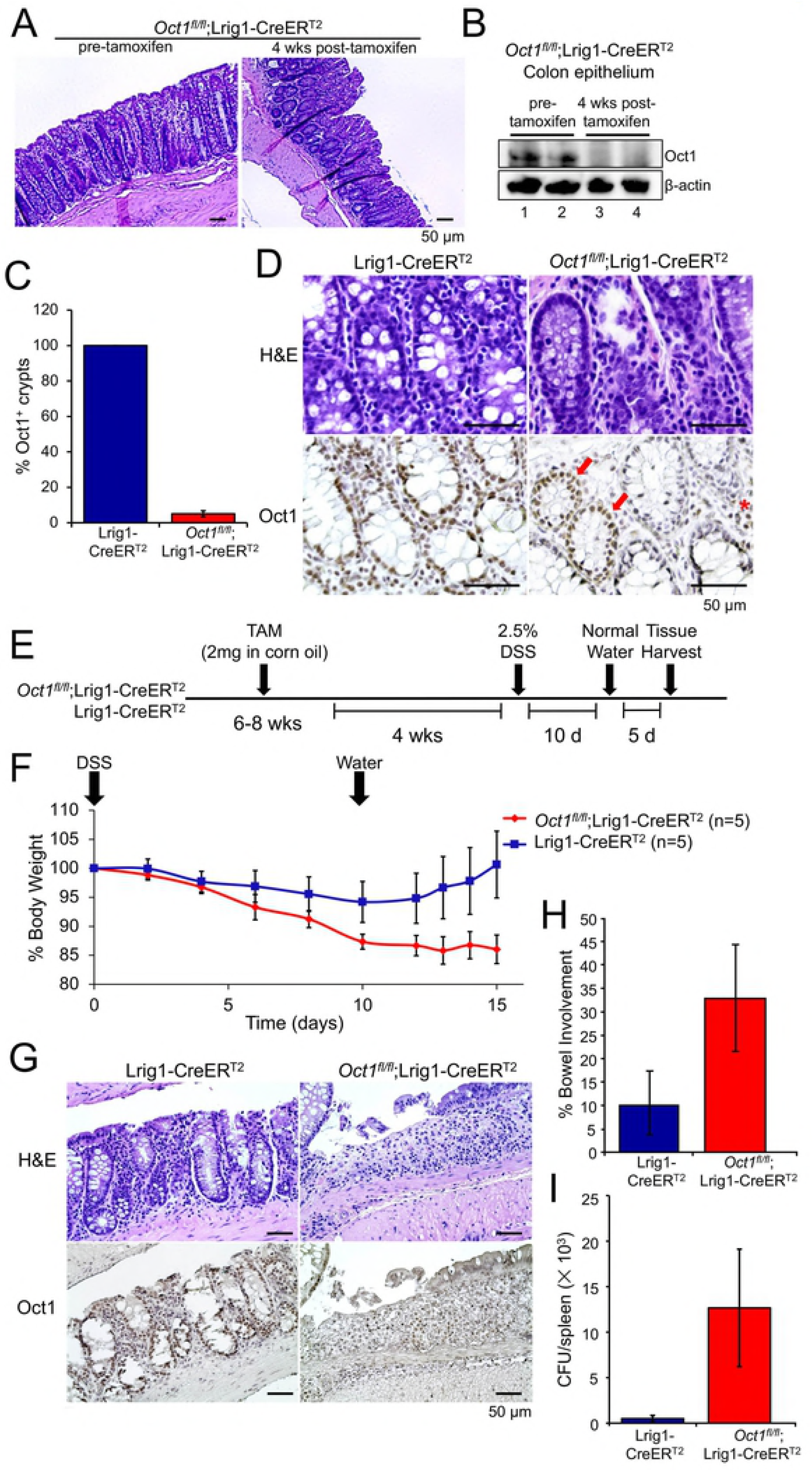
Loss of Oct1 renders mouse colon stem cells more susceptible to DSS-induced damage. Lrig1-CreER^T2^;*Oct1*^*fl/fl*^ mice were treated with tamoxifen to induce the deletion of Oct1 in the Lrig1^+^ cells of the colon. Representative H&E images of the proximal colon are shown. (B) Colon epithelial cells from mice used in (A) were isolated, and subjected to immunoblotting using anti-Oct1 antibodies. β-actin was used as a loading control. (C) Colon sections from mice in (A) were subjected to IHC using Oct1 antibodies. Representative images are shown. Above shows H&E stained adjacent sections of the same tissue. (D) Averages of IHC data from 5 mice are shown. Error bars denote ±standard deviation. (E) Experimental schematic. Lrig1-CreER^T2^;*Oct1*^*fl/fl*^ mice (n=5) and Lrig1-CreER^T2^ controls (n=5) were treated with tamoxifen. 4 weeks post-tamoxifen, mice were treated with 2.5% DSS in their drinking water for 10 days. (F) Average body weight changes of the mice were monitored following DSS treatment. Error bars denote ±SEM. (G) Representative Oct1 IHC images of distal colon sections from mice following 10 day treatment with DSS and 5 day treatment with water. Above shows H&E stained adjacent sections of the same tissue. (H) Images from entire colon (excluding cecum) of 5 mice in each group were evaluated for colitis involvement. Average involvement is shown. Error bars denote ±standard deviation. (H) Spleen homogenates from the same mice as in (H) were grown on blood agar plates to determine bacterial burden and loss of gut barrier function. Average CFUs are shown. N=5 for each group. Error bars denote ±standard deviation.

We treated animals with 2.5% DSS to damage the GI tract and mobilize gut stem cells to regenerate the epithelium (Fig 1E). Mice lacking Oct1 in their colon were somewhat more sensitive to DSS treatment (Fig 1F, day 0-10), consistent with the finding that cells lacking Oct1 are hypersensitive to stress [20]. Upon switching back to water, Oct1 sufficient mice rapidly began to gain weight, while mice lacking Oct1 in the colon continued to lose weight (Fig 1F, day 12-15) and did not regenerate their colon epithelia (Fig 1G). The failure to regenerate was associated with increased colitis (Fig 1H) and loss of barrier function with microbial infiltrate in the spleen (Fig 1I) and liver (data not shown). Collectively, the data indicate that Oct1 loss from the colon results in failure to regenerate following DSS-mediated damage.

### Oct1 is required for regeneration of the small intestinal epithelium following DSS-mediated damage, and required for the passage of intestinal organoids in vitro

To determine if Oct1 behaves similarly in the small intestine, we deleted Oct1 using Lgr5-EGFP-IRES-CreER^T2^ mice [33]. These mice express tamoxifen-inducible Cre and GFP under the control of the native *Lgr5* gene locus, which encodes a R-Spondin receptor expressed in stem cells of the small intestine that stabilizes Frizzled receptors, allowing for increased Wnt signaling [34]. As with the colon, deletion of Oct1 in the small intestine preserved a normal-appearing small intestine (Fig 2A) despite efficient deletion from the epithelium (Fig 2B). Following a similar scheme as in the colon (Fig 2C), failure to thrive and increased weight loss was observed upon removal of DSS from drinking water (Fig 2D). Histological examination revealed that the small intestine had regenerated crypt structures but failed to regenerate nutrient-and fluid-absorbing villi after DSS removal (Fig 2E). These results are similar to the colon in that Oct1 loss results in regeneration-specific defects, but differ in that only villi are affected, with little associated colitis.

**Fig 2.**
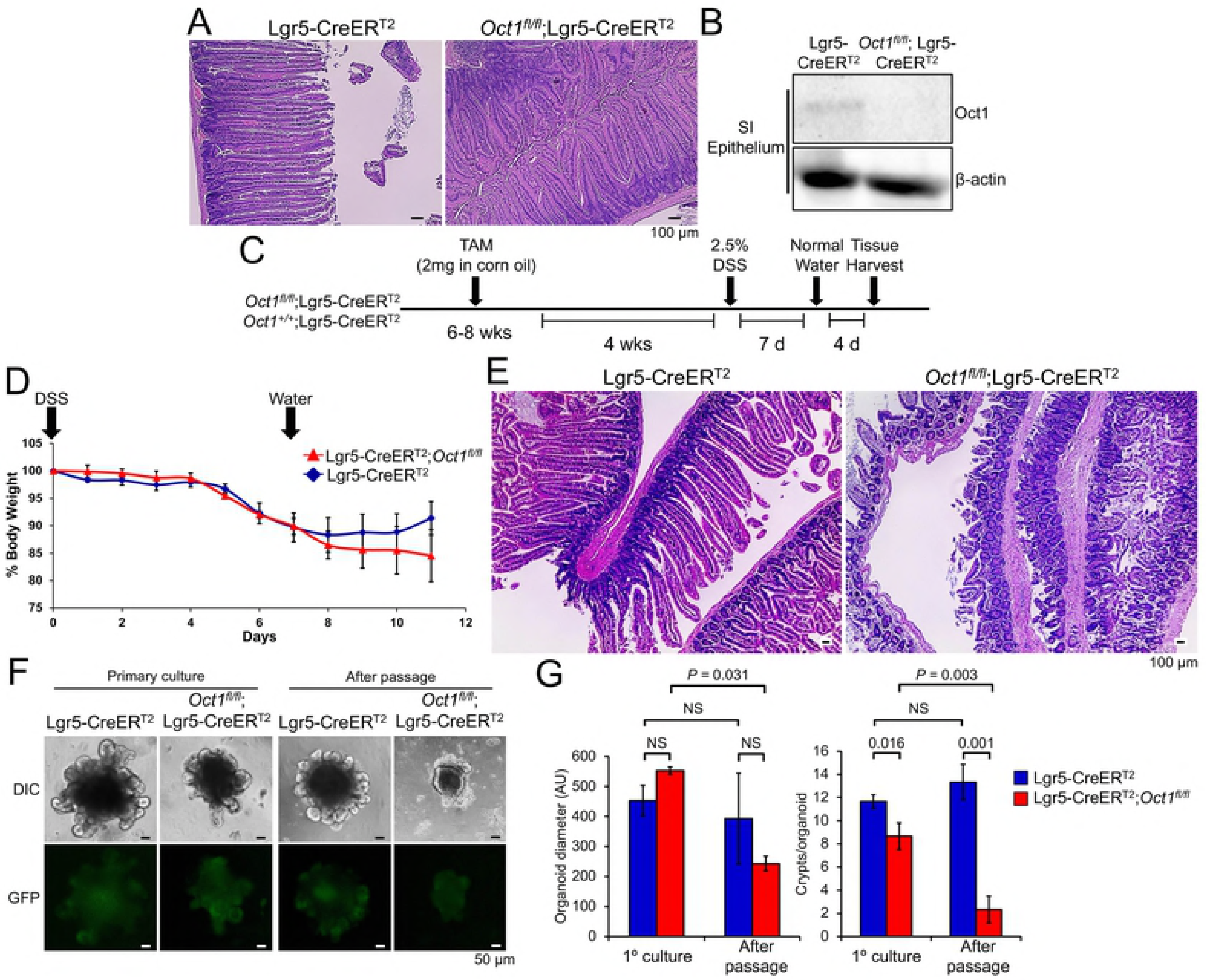
Oct1 is required for regeneration in the small intestine and to maintain gut organoids *in vitro*. (A) Lgr5-EGFP-IRES-CreER^T2^;*Oct1*^*fl/fl*^ mice (n=6) and Lgr5-EGFP-IRES-CreER^T2^ controls (n=4) were treated with tamoxifen for 4 weeks. Representative H&E images of the duodenum are shown. Small intestinal epithelial cells from mice used in (A) were isolated, and subjected to immunoblotting using anti-Oct1 antibodies. β-actin was used as a loading control. (C) Experimental schematic. Lgr5-CreER^T2^;*Oct1*^*fl/fl*^ mice (n=5) and controls lacking the conditional allele (n=5) were treated with tamoxifen. 4 weeks post-tamoxifen, mice were treated with 2.5% DSS in their drinking water for 10 days. (D) Average body weight changes of the mice were monitored following DSS treatment. Error bars denote ±SEM. (E) Representative H&E images of small intestine (duodenum sections from mice following 10 day treatment with DSS and 5 day treatment with water. (F) Tamoxifen-treated mice were used to form intestinal organoids in culture. Images are shown of primary organoids (left) and after one passage (right). (G) Quantification of organoid diameter and average number of crypt domains per organoid from two mice of each genotype (5 organoids/condition/mouse). AU=arbitrary units calculated in Image J.

In vitro cultured intestinal organoids are a powerful tool to study intestinal stem cells and their differentiated progeny. They are relatively easy to culture and manipulate [33,35]. We additionally profiled the effect of Oct1 loss in the small intestine using organoids from mice 4 weeks post-tamoxifen treatment. Organoids were generated directly ex vivo from intestinal crypts. No difference in size, viability or morphological were noted (Fig 2F, primary culture). However, upon passage *Oct1*^*fl/fl*^;Lgr5-EGFP-IRES-CreER^T2^, disaggregated crypts were unable to regenerate new villus and crypt structures to generate complete organoids, resulting in severe abnormalities (Fig 2F, after passage). In contrast, control Lgr5-EGFP-IRES-CreER^T2^ organoids grew normally following passage.

### Oct1 in the colon promotes AOM-DSS-mediated tumorigenicity

Colon cancer mimics a chronically regenerating state in many respects [36,37]. To test if Oct1 loss protects mice from malignancy, we used *Oct1*^*fl/fl*^;Lrig1-CreER^T2^ mice together with a chemical carcinogenesis model driven by the DNA alkylating agent azoxymethane (AOM) [38,39] (Fig 3A). Colon tumors were efficiently generated in control mice but were fewer in number and size in Oct1 deleted mice (Fig 3B). Quantification from 6 experimental and 4 control mice is shown in Fig 3C. In the *Oct1*^*fl/fl*^;Lrig1-CreER^T2^ tumors that did occur, a high proportion of tumor tissue stained positively for Oct1 compared to gross uninvolved (GU) tissue the same section (∼10%, Fig 3D). Quantification from multiple mice indicated that ∼90% of tumor tissue retained Oct1 staining, while only ∼10% of GU tissue did so (Fig 3E). This result indicates a selection against Oct1 deletion in the low-grade tumors that do occur, consistent with Oct1 promotion of tumorigenicity in this model.

**Fig 3.**
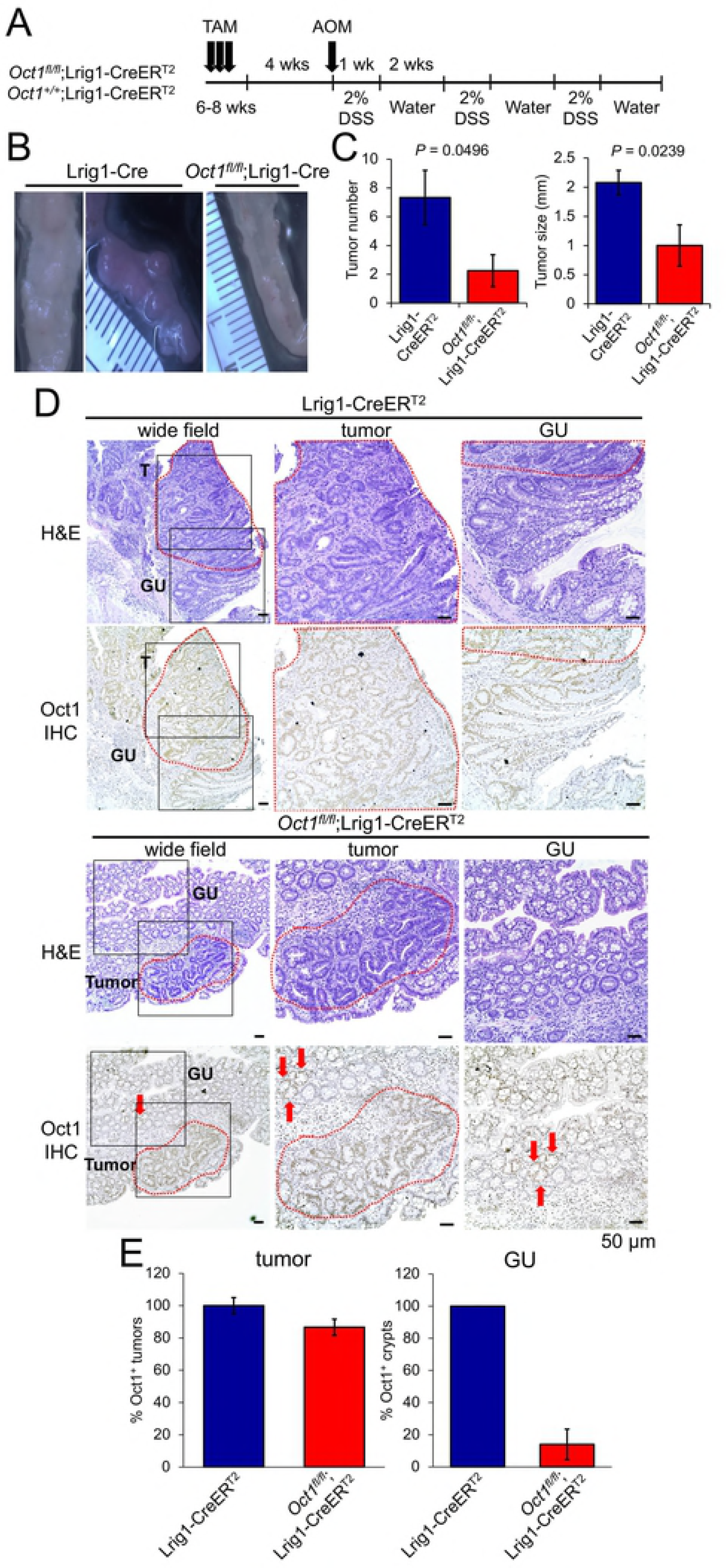
Oct1 loss protects mice from tumor progression in a model of colon cancer induced by AOM/DSS. (A) Lrig1-CreER^T2^;*Oct1*^*fl/fl*^ mice (n=6) and Lrig1-CreERT2 controls (n=4) were treated with tamoxifen followed by a single IP injection of AOM (10mg/kg) 4 weeks later. Mice were then exposed to 3 cycles of 2% DSS in their drinking water for 1 week followed by 2 weeks of water. (B) Images of distal colon tumors from representative mice. (C) Quantification of tumor number and size. (D) H&E and Oct1 IHC of colon tumor tissue and gross uninvolved (GU) adjacent tissue. (E) Analysis of Oct1 IHC in tumor and GU tissue.

### Oct1 restricts tumorigenicity in a model of colon cancer driven by loss of heterozygosity

Oct1 functions physiologically not to promote tumors, but rather to promote stem cell potency. The stem cell properties that Oct1 promotes are largely pro-oncogenic, but in one respect Oct1 can be tumor suppressive: like its paralog Oct4 [40], Oct1 promotes mitotic stability [16], a hallmark of stem cells [40-42]. To test the hypothesis that loss of Oct1 can accelerate tumor initiation in models of malignancy dependent on mitotic errors and LOH, we used conditional deletion of the *Apc* gene, which is mutated in a large proportion of human colon cancers [43]. Over time, *Apc* LOH results in adenocarcinomas in the distal colon that mimic human disease in many respects [32]. We crossed *Apc*^*fl*^ mice [44] with *Oct1* (*Pou2f1*) conditional mice, generating an allelic series of *Oct1*^*+/+*^, *Oct1*^*+/fl*^, and *Oct1*^*fl/fl*^ Lrig1-CreER^T2^ mice with heterozygous *Apc*^*fl*^. Mice were followed for 100 days post-tamoxifen treatment before sacrifice. Progressive deletion of one or both *Oct1* alleles progressively increased gross tumor number in this model (Fig 4A). Quantification of H&E sections from multiple animals confirmed that tumor number was increased (Fig 4B) but average area per tumor was not (Fig 4C). Using Oct1 IHC we also found that Oct1 was again efficiently deleted from normal (gross uninvolved) crypts, with ∼10% of crypts escaping deletion (Fig 4D, arrows). In this case and in contrast to AOM-DSS-mediated tumors, Oct1 was deleted in most tumor cells (Fig 4D). Oct1 was present in non-epithelial cells within the tumor, providing a measure of specificity (Fig 4D, asterisks). We also assessed β-catenin by IHC. Apc protein restricts β-catenin by ubiquitin-mediated degradation [45], and hence *Apc* LOH would be predicted to augment β-catenin specifically in tumors in this model. As expected, we found accumulated β-catenin, including nuclear β-catenin, in the tumor cells (Fig 4D).

**Fig 4.**
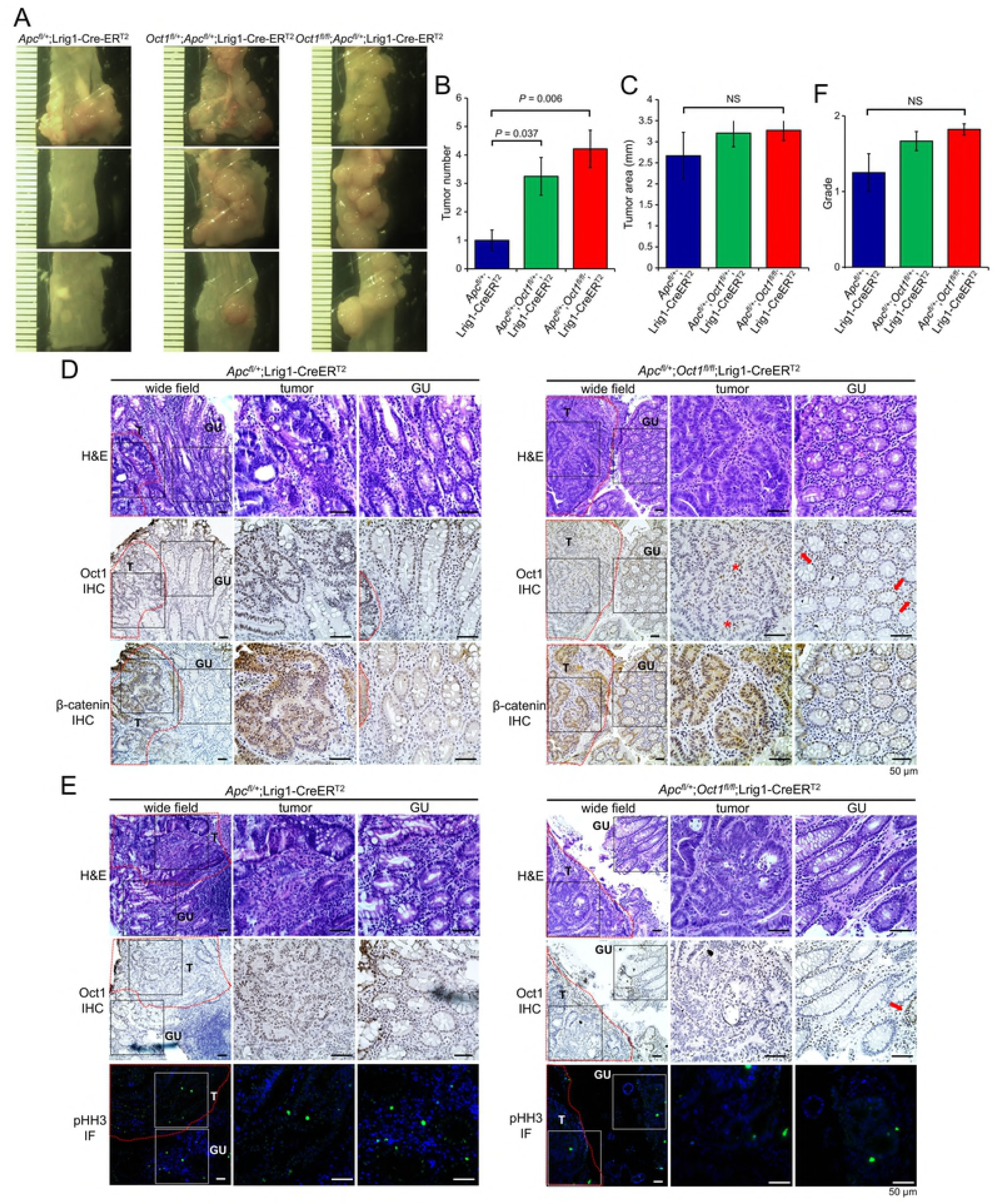
Oct1 restricts tumorigenicity in a model driven by heterozygous *Apc* deletion in Lrig1^+^ cells. (A) Example distal colons are shown of an allelic series of *Oct1*^*+/+*^, *Oct1*^*fl/+*^ and *Oct1*^*fl/fl*^ animals that were additionally *Apc*^*fl/+*^ and Lrig1-CreER^T2^. (B) Quantification of tumor incidence across multiple animals. N=5 for each group. Error bars denote ±standard deviation. (C) Similar analysis performed for tumor size. (D) Representative H&E, and Oct1 and β-catenin IHC of colon tumor tissue and adjacent GU tissue from *Apc*^*fl/+*^;Lrig1-CreER^T2^ and *Apc*^*fl/+*^;*Oct1*^*fl/fl*^;Lrig1-CreER^T2^ animals. Example tumor stromal cells with Oct1 staining are shown with the asterisk. Example normal crypt structures in GU tissue that escape Oct1 deletion in *Apc*^*fl/+*^;*Oct1*^*fl/fl*^;Lrig1-CreER^T2^ animals are shown with arrows. Mice were analyzed 100 days after tamoxifen treatment and *Apc*/*Oct1* deletion. (E) Tumor mitoses were studied using phospho-histone H3 IF. H&E and Oct1 IHC are shown for comparison. Arrow indicates position of example crypt in GU tissue that escaped Oct1 deletion in *Apc*^*fl/+*^;*Oct1*^*fl/fl*^;Lrig1-CreER^T2^. (F) Quantification of tumor grade across multiple animals. N=5 for each group. Tumor grade was low in all cases. Error bars denote ±standard deviation.

To study tumor aggressiveness in this model, we conducted immunofluorescence (IF) using phospho-histone H3 antibodies. The number of mitotic events was low, and equivalent in the presence or absence of Oct1 (Fig 4E). Consistent with this result, pathological scoring of tumor sections indicated that despite the increased tumor incidence, there was no significant difference in tumor grade (Fig 4F).

### Genes that are differentially expressed between chemical and Apc-LOH tumors are preferentially enriched for Oct1 targets

The finding that Oct1 loss resulted in opposing effects in the chemical-vs. *Apc*/LOH-driven colon cancer models suggested that the two models may differ at a molecular level, such that Oct1’s dominant activity can switch from pro-oncogenic to tumor suppressive. To test this hypothesis, we sampled gene expression from control Oct1 wild-type FFPE and frozen tumor samples. We used a custom 60-gene panel enriched in gut-associated stem cell function together with 31 AOM-DSS-induced and 25 *Apc*^*fl*^;Lrig1-CreER^T2^ samples, all of which were wild-type for Oct1. Unsupervised hierarchical clustering of gene expression resulted in interdigitation of the samples (Fig 5A, top). The interdigitation was robust using multiple settings and cutoffs (not shown), indicating that global tumor-to-tumor gene expression variation is dominant over differences between the models. To identify model-associated molecular signatures, we grouped samples by the model of origin and identified subsets of profiled genes whose expression partitioned with the model. A group of 17 genes was identified (Fig 5A, bottom) whose expression tended to correlate with tumor model (Fig 5B, Table S1). We compared this group of genes with Oct1 targets identified in T cells [18] and with Oct1/Oct4 targets identified in embryonic stem cells (ESCs, [46]. Oct4 levels are far higher than Oct1 in ESCs, resulting in most targets being occupied exclusively by Oct4, which has similar DNA binding specificity [46]. Of the 17 identified genes whose expression correlates with the tumor model, 8 were previously identified as direct Oct1 targets in T cells and 9 as Oct4 targets in ESCs (Fig 5C). Additionally *Lef1*, a gene on the tumor panel expressed more strongly in the *Apc*-LOH-driven model (Fig 5B), was identified as one of only 356 direct Oct1-exclusive target genes in ESCs. Across both ChIPseq datasets, all but four of the 17 genes overlapped with direct Oct1 targets (Table S1). To further test the robustness of this result, we performed a second analysis with a larger 770 gene panel together with 23 AOM-DSS-induced and 12 *Apc*^*fl*^;Lrig1-CreER^T2^ Oct1 wild-type samples, in this case identifying 83 differentially expressed genes (Table S1). These genes were also enriched for Oct1 and Oct4 targets (Fig 5C). In this case, three of the genes overlapped with a set of Oct1-exclusive ESC targets: *Ercc2, Lifr* and *Myb* (Table S1). The enrichment for Oct1/Oct4 target genes in the differentially expressed gene set was statistically significant in all cases (Fig 5C). Collectively, these results indicate that Oct1 regulates genes that tend to be differentially expressed tumors from the two models, potentially contributing to the opposing effects of Oct1 in the two models.

**Fig 5.**
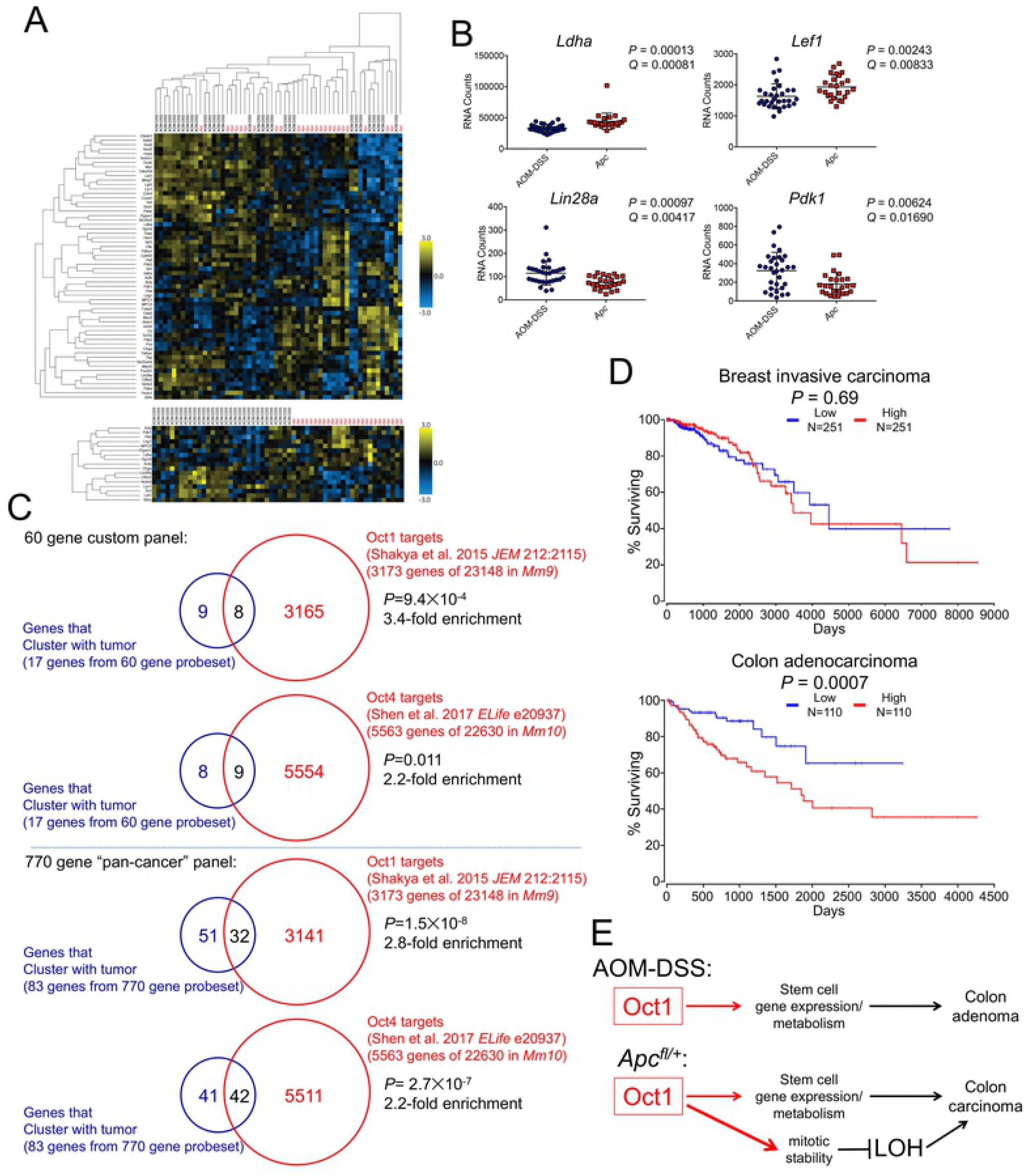
Effect of *Oct1* mRNA expression level in human colon and breast cancer, and model for Oct1 function in different mouse models of gut malignancy. (A) Gene expression analysis performed using a custom probeset with 31 AOM-DSS tumors (23 FFPE, 8 snap-frozen) and 25 *Apc*^*fl/+*^-LOH tumors (12 FFPE, 13 snap-frozen). Top: A heatmap is shown using unsupervised hierarchical clustering using all genes in the probeset. Bottom: Forced clustering by tumor type, showing top clustering of genes based upon mean change and ranked adjusted *P*-value. (B) Example genes with significant expression differences based on tumor type. (C) Intersection of genes enriched from custom probeset with Oct1 and Oct4 targets previously identified by ChIPseq in other cell types. A second analysis is also shown using genes enriched from a larger PanCancer Pathways probeset (Table S1). (D) Kaplan-Meier survival curves were downloaded and modified from the OncoLnc site (http://www.oncolnc.org/, gene symbol: POU2F1), and compare the top and bottom quartiles of human breast and colon cancer patients in each plot, respectively. (E) Functions of Oct1 are shown in AOM/DSS colon tumor model and *Apc*^*fl/+*^;Lrig1-CreER^T2^ colon tumor model. In the case of *Apc*^*fl/+*^ tumors based on LOH, Oct1 gains a tumor suppressive activity by promoting mitotic stability.

## Discussion

Here we show that the transcription factor Oct1 is dispensable for mouse gut epithelial homeostasis, but is essential for regeneration and the passage of intestinal organoids in vitro. Oct1 can be deleted in the colon with normal architecture for up to 150 days, consistent with the interpretation that Oct1 is dispensable for maintenance of the normal gut. In contrast to homeostatic conditions, Oct1 is required to regenerate the gut following DSS exposure. The findings are consistent with prior findings from others indicating that homeostasis and regeneration are molecularly and physiologically distinct [47-51]. DSS treatment of mice lacking Oct1 in small intestinal epithelium also had no effect on homeostasis, but resulted in continued weight loss after switching back to water. However the effect was weaker than in the colon and inflammation and loss of barrier function were not observed. The difference is likely due to the fact that crypts but not fluid-and nutrient-absorbing villi were efficiently generated. The difference in regeneration potential between the two organs and Cre drivers (Lrig1 vs. Lgr5) is unknown.

Intestinal organoids have been used before to study regeneration [52]. Consistent with the finding that Oct1 loss did not affect small intestinal homeostasis in vivo, organoids could be directly explanted from tamoxifen treated mice. Interestingly, we found that intact organoid structures could not be maintained by passage in vitro. Instead isolated crypts structures could close but not generate new villus and crypt domains. The underlying causes may be similar to the failure to regenerate in vivo, though more study is required to test this idea.

We also show that Oct1 loss has potent effects on tumorigenicity in two different mouse models of colon malignancy. In one model (AOM-DSS), Oct1 loss strongly protects mice from tumors. Elevated Oct1 mRNA expression is a negative prognostic factor in colon but not breast cancer (Fig 4D). The normal appearance of the colon following Oct1 deletion, coupled with the protection afforded by Oct1 loss, suggests a possible “therapeutic window” in which targeting Oct1-associated pathways could be used to treat GI malignancies with minimal side effects. However, in a second model (*Apc*^*fl/+*^;Lrig1-Cre), Oct1’s dominant activity is tumor suppressive, with more tumors generated though of equal grade. More study will be required to determine the conditions in which targeting Oct1 and its associated pathways in familial and non-familial gut malignancy could be beneficial.

In HeLa cells and MEFs, Oct1 loss slightly increases the rate of mitotic chromosome segregation abnormalities, resulting in increased lagging chromosomes and aneuploidy [16]. Consistent with these prior findings, Oct1 loss accelerated tumorigenesis and increased tumor number in a colon cancer model driven by LOH of the tumor suppressor gene *Apc* in Lrig1^+^ cells [32]. In additions to increased LOH, differences in the molecular pathways and vulnerabilities associated with Oct1 in the two tumor models could contribute to the difference. To test this idea, we profiled gene expression in AOM-DSS tumors and tumors in which *Apc* is deleted in Lrig1^+^ cells, identifying a set of differentially expressed genes. These genes were enriched in Oct1 targets. The results were robust, as they were reproduced with two different gene expression panels. Differential regulation of gene expression by Oct1 in the two models could therefore explain in part the opposing effects of Oct1 loss in the two models. A model for the effect of Oct1 loss in the two tumor models is shown in Fig 4E. In this model, Oct1 promotes AOM/DSS-induced tumors through actions on target genes controlling metabolism and stem cell identity. Because Oct1 promotes mitotic stability and because Oct1 target genes are differentially expressed in the more aggressive *Apc*-LOH model, Oct1 instead acts as a tumor suppressor. Cumulatively, the findings indicate that Oct1 is a potent regulator of colon malignancy, but that its functions are dictated by the colon tumor model used.

## Materials and methods

### Laboratory mice

All mice used in this study were mixed C57BL/6:129/Sv background. The Oct1 conditional allele has been described previously [18]. The Lrig1-CreER^T2^ mouse allele [32] was a gift of Robert Coffey (Vanderbilt). The Lgr5-EGFP-IRES-CreER^T2^ allele [33] was purchased from Jackson labs. *Apc*^*loxp exon14*^ *(Apc*^*fl*^) has been previously described [44] and was a gift from Ömer Yilmaz (MIT). Food and water were available *ad libitum*. Tamoxifen (200 µL 10 mg/mL in corn oil) was administered by oral gavage. All mice were treated at 6-8 weeks of age. Regeneration and azoxymethane (AOM) chemical tumorigenesis experiments used a single tamoxifen treatment. The *Apc* tumor model received three treatments on consecutive days. Dextran sodium sulfate (DSS, MP Biomedicals) was provided in drinking water at a concentration of 2.0% for AOM-induced tumors, and 2.5% for regeneration. AOM (Sigma) was provided by IP injection (10mg/kg). AOM-DSS treatments followed the protocol in [39]. All *in vivo* experiments were reviewed, approved, and conducted in compliance with the University of Utah’s Institutional Animal Care and Use Committee and the NIH Guide for Care and Use of Laboratory Animals guidelines.

### Immunoblotting

Antibodies used for immunoblots were as follows: Oct1, Bethyl #A310-610; β-actin, Santa Cruz #sc-47778.

### Preparation of colon and small intestine epithelial cells and crypts

Total epithelial cells from the colon were isolated as previously described [53,54].

### Immunohistochemistry

IHC was performed as in [7]. The slides were developed with DAB peroxidase substrate (Vector Laboratories, SK-4100) as per manufacturer instructions, and were counterstained with hematoxylin. After dehydration (3 min washes each of 70%, 85%, 95% and 100% ethanol) the slides were incubated in xylene for 3 min twice and mounted using Limonene mounting medium (Abcam #104141). Antibodies used for IHC were as follows: Oct1, Abcam #178869 and ß-catenin, Cell Signaling #8814.

### Immunofluoresence

IF was performed as in [7] using an antibody against phospho-histone H3 (Abcam #5176). The slides were mounted using Fluoroshield mounting medium with DAPI (Abcam #104139).

### Intestinal organoids

Organoids were maintained as previously described [53,54] with modifications. Crypts from tamoxifen-treated or untreated mice were plated in 8-well chambered slides in 40µl of Matrigel at a density of ∼40 crypts per Matrigel droplet. Organoids were grown for 5-7 days until fully grown. Mature organoids were passaged every 5-7 days. Images were taken using an Olympus FV1000 confocal inverted microscope. Quantifications were performed using Image J software (National Institutes of Health).

### Tumor gene expression profiling

Gene expression was measured in Oct1 wild-type formalin-fixed, paraffin-embedded (FFPE) and snap-frozen tumors using NanoString (Seattle, WA). For frozen samples, tumors were dissected from the normal mucosa under an Olympus SZ61 dissecting microscope in ice cold PBS and snap frozen in liquid nitrogen. Two gene panels were used: the 770-gene “pan-cancer” panel (23 FFPE AOM-DSS tumors and 12 FFPE *Apc*^*fl/+*^;Lrig1-CreER^T2^ tumors), and a custom 60 gene panel enriched for gut stem cell-associated genes [54] (23 FFPE and 8 frozen AOM-DSS tumors, and 12 FFPE and 13 frozen *Apc*^*fl/+*^;Lrig1-CreER^T2^ tumors). Laser capture microdissection was performed by the Molecular Pathology Core at the University of Utah using a Qiagen miRNeasy FFPE Kit (217504). Analysis and normalization of the raw data were conducted with nSolver Analysis Software v4.0 (NanoString Technologies). Genes whose expression significantly correlated with tumor type were identified using Prism GraphPad Row Statistics. Multiple T-tests were performed on each gene individually, without assuming a consistent standard deviation. Cutoffs were determined using the Two-stage linear step-up procedure of Benjamini, Krieger and Yekutieli, with *Q* = 1% or 5%. For each differentially expressed gene set, ascription of statistical significance for Oct1/Oct4 target gene enrichment was made using a hypergeometric test.

## Acknowledgements

We thank C. Murtaugh and H. Zhao for critical reading of the manuscript. R. Coffey provided Lrig1-CreER^T2^ mice and Ö. Yilmaz provided *Apc*^*fl*^ mice. We thank J. Round and members of her laboratory for the use of equipment, and J. Gertz for help with statistical analysis. We thank J. O’Shea and members of the University of Utah Health Sciences Center Molecular Diagnostics Core facility for assistance with NanoString.

## Supporting information legends

**Table S1. Genes with expression correlating to tumor model (AOM-DSS vs Apc-LOH;Lrig1-CreER).**

